# MADS-box genes galore in wheat genome: phylogenomics, evolution and stress associated functions

**DOI:** 10.1101/2020.10.23.351635

**Authors:** Qasim Raza, Awais Riaz, Rana Muhammad Atif, Babar Hussain, Zulfiqar Ali, Hikmet Budak

**Affiliations:** Molecular Breeding Laboratory, Rice Research Institute, Kala Shah Kaku, Sheikhupura, Punjab, Pakistan; Department of Plant Breeding and Genetics, University of Agriculture Faisalabad, Pakistan; Center for Advanced Studies in Agriculture and Food Security (CAS-AFS), University of Agriculture Faisalabad, Pakistan; Department of Molecular Biology & Genetics, Middle East Technical University, Ankara, Turkey; Faculty of Life Sciences, University of Central Punjab, Lahore, Pakistan; Institute of Plant Breeding and Biotechnology, Muhammad Nawaz Shareef University of Agriculture, Multan, Pakistan; Montana BioAgriculture, Inc., Bozeman, MT, United States

**Keywords:** Genome-wide analysis, *In-silico* expression, *TaMADS’s*, Transcription factors, *Triticum aestivum*, Wheat adaptability

## Abstract

MADS-box gene family members play multifarious roles in regulating the growth and development of crop plants and hold enormous promise for bolstering grain yield potential under changing global environments. Bread wheat (*Triticum aestivum* L.) is a key stable food crop around the globe. Until now, the available information concerning MADS-box genes in the wheat genome has been insufficient. However, a comprehensive genome-wide analysis identified 300 high confidence MADS-box genes from the latest publicly available reference genome of wheat. Comparative phylogenetic analyses with *Arabidopsis* and rice MADS-box genes classified the wheat genes into 16 distinct subfamilies, without a single *FLOWERING LOCUS C* homolog present in the wheat genome. Gene duplications were mainly identified in subfamilies containing unbalanced homeologs, pointing towards a potential mechanism for gene family expansion. Moreover, a more recent evolutionary origin was inferred for M-type genes, as compared with MIKC-type genes, indicating their significance in understanding the evolutionary history of the wheat genome. We speculate that subfamily-specific distal telomeric duplications in unbalanced homeologs facilitate the rapid adaptation of wheat to changing environments. Furthermore, our *in-silico* expression data strongly proposed MADS-box genes as active guardians of plants against pathogen insurgency and harsh environmental conditions. In conclusion, we provide an entire complement of MADS-box genes identified in the wheat genome that will accelerate functional genomics efforts and possibly facilitate bridging gaps between genotype-to-phenotype relationships through fine-tuning of agronomically important traits.

## 1. Introduction

In the fight for global food security and safety, bread wheat represents one of the largest contributing grain crops. However, following the green revolution, improvement of wheat grain production has been hindered by various bottlenecks. This includes, but is not limited to, non-availability of a reliable and fully annotated reference genome (Brenchley et al., 2012; Chapman et al., 2015; Clavijo et al., 2017; IWGSC, 2014; Zimin et al., 2017). The slow progress in harnessing a fully annotated reference genome was mainly due to the genome’s allohexaploid nature, which comprises three closely related, but independently maintained, sub-genomes, known as A, B, and D. This, in combination with a high frequency of repetitive sequences, particularly hindered progress. Recently, an alliance of geneticists combined resources over a period of 13 years, to produce a fully annotated sequence of the wheat genome, which had a resultant size of ~17 Gbps. This is the largest known genome in crop plants (IWGSC, 2018). To add to this, the high-quality of the reference genome, together with large scale RNA-seq data and expression repositories (Borrill et al., 2016; Ramírez-González et al., 2018), provide a rich resource for studying evolutionary dynamics and functional characterization of important gene families, which in turn could facilitate crop improvement efforts.

MADS-box transcription factors (TFs) are a well-documented group of genes known for playing vital roles in regulating the growth and development of several important plant species (Ali et al., 2019; Smaczniak et al., 2012). These genes influence diverse biological functions, including cell development, signal transduction, biotic and abiotic stress responses, vegetative organs development, control of flowering and anthesis time, formation of meristems and flower organs, ovule development, embryo development, dehiscence zone formation, and ripening of fruits and seeds (Ali et al., 2019; Alvarez-Buylla et al., 2000; Colombo et al., 1995; Kuo et al., 1997; Messenguy and Dubois, 2003; Moore et al., 2002; Pařenicová et al., 2003; Riechmann and Meyerowitz, 1997; Rounsley et al., 1995; Saedler et al., 2001; Samach et al., 2000; Smaczniak et al., 2012). Given their importance, identification and characterization of MADS-box genes in agriculturally important species is critical for crop improvement and fine-tuning of specific-traits through genetic exploitations.

All MADS-box genes contain a highly conserved MADS (M) domain of approximately 58–60 amino acids. The M domain enables DNA binding and is located in the N-terminal region of the protein (Yanofsky et al., 1990). Members of this gene family are classified into M/type I and MIKC/type II super clades, which were generated after an ancient gene duplication event which occurred before the separation of animal and plant lineages (Becker and Theißen, 2003). M-type genes have simple intron-exon structures (zero or one intron), only contain a single M domain, and encode Serum Response Factor (SRF)-like proteins. These can be subcategorized into four clades (*Mα, Mβ, Mγ* and *Mδ*). However, the *Mδ* clade genes closely resemble those within the *MIKC^*^* group, as previously reported (De Bodt et al., 2003).

In comparison, MIKC-type genes have four domains: (i) a highly conserved DNA binding M domain, (ii) a less conserved Intervening (I) domain of ~30 AA involved in dimer formation, (iii) a moderately conserved Keratin (K) domain of ~70 AA which regulate heterodimerization of MADS proteins, and lastly (iv) a highly inconstant C-terminal region which contributes to transcriptional regulation and higher-order protein complex formation (Díaz-Riquelme et al., 2009; Henschel et al., 2002). MIKC-type genes encode Myocyte Enhancer Factor 2 (MEF2)-like proteins and are categorized into *MIKC^c^* and *MIKC^*^* clades (Kaufmann et al., 2005). *MIKC^*^*-type genes have duplicated K domains and relatively longer I domains than *MIKC^c^* -types (Duan et al., 2014). *MIKC^c^*-type genes can be further categorized into several subclades based on their phylogenetic relationships in flowering plants (Gramzow and Theißen, 2015).

Similar to what has been observed in other model and crop plant species, in wheat MADS-box genes are known to confer drought tolerance through regulation of drought tolerance genes and micro-RNAs (Budak et al., 2015). In addition to this, *TaMADS51, TaMADS4, TaMADS5, TaMADS6*, and *TaMADS18* all showed up-regulation, where *TaMADAGL17, TaMADAGL2, TaMADWM31C*, and *TaMADS14* were downregulated, under phosphorous (P) starvation, and further functional analysis confirmed their role in P-deficient stress responses (SHI et al., 2016). In another study, several MADS-box genes were activated and differentially expressed after inoculation of wheat spikes with Fusarium head blight (FHB) (Kugler et al., 2013). Furthermore, a simple yet elegant ABCDE floral organ identity model explained the complex genetic interactions among MIKC-type wheat MADS-box genes for determining the fate of floral organs (Ali et al., 2019). These significant roles of MADS-box genes in biotic and abiotic stresses, fertilizer response, and flower development highlight the necessity of their comprehensive identification and characterization in the bread wheat genome.

Previously, comprehensive genome-wide analyses of MADS-box genes have been carried out in diverse plant species. However, the available information about wheat MADS-box genes is comparatively sparse. For example, previously reported genome-wide identification of MADS-box genes in wheat (Ma et al., 2017; Schilling et al., 2020) has been revealed as outdated and inadequate, as it was performed by Ma et al. (2017) using only a draft version of the wheat genome (TGACv1) which contains a lesser number of genes. To add to this, Schilling et al. (2020) studied only MIKC-type MADS-box genes. Moreover, a significant number of low confidence and/or truncated protein-coding pseudogenes were also included in the before mentioned studies.

In order to bridge the gaps in knowledge, in this study we comprehensively identify the entire complement of high confidence, full-length, protein-coding MADS-box genes present in the latest publicly available reference genome of bread wheat (IWGSC RefSeq v1.1). Through a combination of different search approaches, 300 high confidence, non-redundant, full-length MADS-box genes were identified. Comparative phylogenetic analyses with model plant MADS-box genes further classified these into at least 16 *Arabidopsis* and/or grass specific subfamilies. Moreover, homeologous and duplicated genes were identified to study probable gene family expansion and evolution mechanisms. Furthermore, expression patterns of identified MADS-box genes were studied under several biotic and abiotic stress conditions. In this way, we provide a comprehensive resource of wheat MADS-box genes which have the potential to facilitate molecular breeders in fine-tuning of important traits for further improvement.

## 2. Results

### 2.1. MADS-box genes galore in wheat genome

In this study, conserved domains and query search-based approaches identified a total of 300 high confidence and non-redundant MADS-box genes from the latest publicly available wheat genome (**Table S2**). The NCBI CDD batch search further revealed that 167 (~55.7%) encode MADS-box proteins, 125 (~41.7%) encode MADS and K-box proteins, and only 8 (~2.6%) encode K-box domain-containing proteins (**Table S3**). The proteins lengths, molecular weight, and isoelectric points of MADS domain-containing proteins ranged from 58–450 amino acids, 6.483–48.304 kD, and 4.420–12.123 pI, respectively (**Table S2**). This data suggests that different MADS-box genes may function within different environments.

Comparative phylogenetic analyses between *Arabidopsis* and wheat MADS-box genes (Fig. S1), as well as between rice and wheat (Fig. S2), distinguished wheat genes into M- and MIKC-types (**Table S2**). In wheat, 128 (~43%) MADS-box genes exhibited high sequence similarity with *Arabidopsis* and rice M-type (type-I) genes, whereas 172 (~57%) revealed more homology with MIKC-type (type-II) genes (Fig. 1). In general, MADS-box genes were equally distributed among the 21 wheat chromosomes, with an exception of the three homeologous chromosomes (7A, 7B and 7D), which harboured a significantly higher number of genes and displayed peak gene density values (Fig. S3). However, M-type genes were randomly distributed among chromosomes and were predominantly located on three homeologous chromosomes of 3^rd^, 6^th^, and 7^th^ linkage groups (Fig. S4). In comparison, MIKC-type genes were equally distributed on all chromosomes. Furthermore, the percent contributions of A, B, and D genomes were also comparable (Fig. S3). As expected, MIKC-type genes were the largest group of MADS-box genes in wheat. To date, the wheat genome harbours the 2^nd^ largest number of MADS-box genes identified, following *Brassica napus* (Wu et al., 2018). Possible explanations for the abundance in wheat includes the larger genome size, high gene number, high rate of homeolog retention, and hexaploid nature of the genome.

**Fig. 1.**
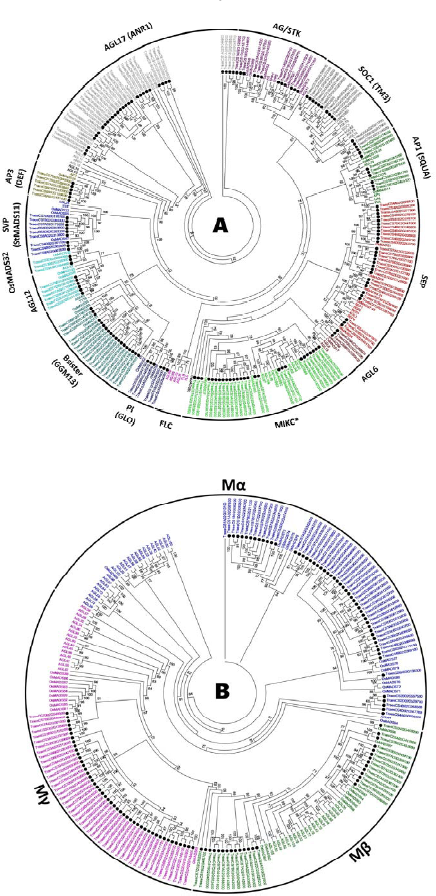
Comparative phylogenetic analysis based subfamily classifications of wheat MADS-box genes. *Arabidopsis*, rice and wheat MADS-box proteins were MAFFT aligned (Katoh et al., 2018; Katoh and Standley, 2013), maximum likelihood phylogenies inferred using I_Q_-T_REE_ software (Nguyen et al., 2015) and final trees visualized with MEGA7 (Kumar et al., 2016). Separate phylogenetic trees were inferred among Arabidopsis, rice and wheat MIKC-type (**A**) and M-type (**B**) proteins. Wheat MADS-box genes are highlighted with solid black circles. Only bootstrap values ≥50%, as calculated from 1000 replicates, could be displayed on the tree nodes. Subfamily-specific colouring was adopted for differentiating the different subfamilies. Subfamily names (outer bands) were given by following the subfamily classifications in Arabidopsis and/ or major grass species (Gramzow and Theißen, 2015).

### 2.2. Subfamily diversity in MADS-box genes

Separate ML phylogenies among *Arabidopsis*, rice, and wheat MADS-box genes exposed 14 MIKC-type and 3 M-type major subfamilies. The 172 MIKC-type genes were unevenly dispersed into *AP1* (9), *AP3* (6), *PI* (6), *AG*/*STK* (12), *SEP* (28), *AGL6* (3), *AGL12* (6), *AGL17* (31), *Bsister* (19), *MIKC^*^* (27), *OsMADS32* (3), *SOC1* (13), and *SVP* (9) subfamilies (Fig. 1A, **Table S2**). As expected, monocot (*OsMADS32*) and eudicot (*FLC*) specific gene subfamilies were also witnessed. Absence of monocot orthologs from eudicot specific subclade and/or eudicot orthologs from monocot specific subclade suggest that genes of these subclades might have lost after the divergence of monocot and eudicot lineages. Likewise, 128 M-type genes were randomly distributed into *Mα* (53), *Mβ* (28), and *Mγ* (47) subfamilies (Fig. 1B).

In general, *Arabidopsis*, rice and wheat MADS-box gene subfamilies roughly followed the species-specific phylogenetic clades. Triads of wheat homeologous genes exhibited close relationships with one or more rice genes, with *Arabidopsis* genes representing a sister group association to grass genes (e.g. the *AP1, AP3, PI, AG/STK, AGL6, AGL12, OsMADS32, SOC1*, and *SVP* subfamilies; Fig. 1). Whereas, the subfamily phylogenies were more complex in case of *AGL17, Bsister*, MIKC^*^, *SEP*, and all M-type MADS-box genes, probably due to multiple duplication events during the polyploidization of wheat genome.

### 2.3. Wheat genome lacks FLC-like genes

*FLC-like* genes have been confirmed to regulate flowering time and vernalization responses in plants (Andrés and Coupland, 2012; Distelfeld et al., 2009) and their wheat and rice counterparts have previously been reported (Ruelens et al., 2013; Schilling et al., 2020). However, in this study, despite employing updated resources and tools (see Materials and Method section), none of the wheat and rice MADS-box genes fell into the *Arabidopsis* specific *FLC*-clade (Fig. 1A). All the wheat genes which were previously reported to be *FLC*-like (**Table S4**) were grouped with MIKC^*^-like MADS-box genes. Furthermore, we demonstrated that amino acid sequences of *Arabidopsis* specific *FLC* genes were significantly different from MIKC^*^-like genes (Fig. 2). The most significant differences were detected in MADS domain region at 30^th^, 34^th^, and 50^th^ positions where glutamic acid (E), glutamine (Q), and alanine/glycine/serine (A/G/S) residues of *FLC* genes were substituted with lysine (K), glutamic acid (E), and proline (P), respectively. Collectively, these results strongly indicate that wheat and rice genomes lack *FLC* genes, which might have been lost after the divergence of eudicot and grass lineages.

**Fig. 2.**
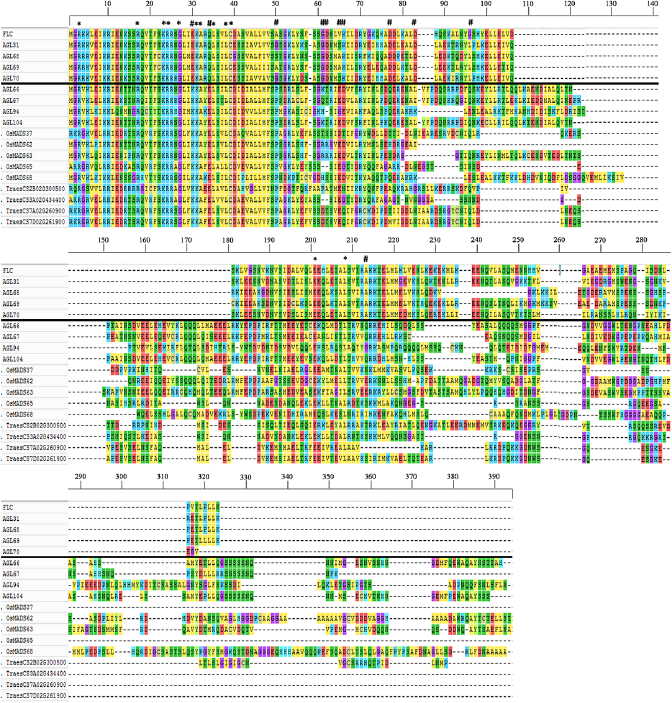
Structural differentiation between *FLC*-like and *MIKC^*^*-like MADS-box genes. Amino acid sequences of representative genes from both subfamilies were aligned using Clustal Omega (Sievers et al., 2011) with default parameters and multiple sequence alignment visualized with MEGA7 (Kumar et al., 2016). *FLC*-like genes (above solid black line) were separated from *MIKC^*^*-like genes (below solid black line). The conserved and diverged residues were indicated with a star (*) and asterisk (#) symbols, respectively. Multiple sequence alignment demonstrated conservation of amino acid residues/motifs within both subfamilies, whereas diversification between subfamilies.

### 2.4. Subfamily-specific gene duplications in MADS-box genes

Overall, MADS-box genes were equally located in the interstitial and proximal regions (R2a, R2b and C) and distal telomeric (R1 and R3) ends of chromosomes (49% and 51%, respectively) (**Table S2**). However, substantial differences were observed among gene locations of M-type and MIKC-type genes, as well as among subfamilies. The majority of the *M*-type genes were located in distal telomeric segments (62%), whereas MIKC-type genes were more prevalent in central chromosomal segments (57%). Generally, a larger portion of genes belonging to significantly expended subfamilies tended to be located in distal telomeric ends, whereas genes of smaller subfamilies were more clustered in central chromosomal segments (**Table S2**).

Gene duplications were identified through sequence similarities in coding sequences of all MADS-box genes. A total of 201 duplicated gene pairs with ≥ 90% sequence homology were identified, which corresponded to 123 non-redundant genes (Fig. 3; **Table S6**). Two genes (*TraesCSU02G209900, TraesCSU02G235300*) with unknown chromosomal location information were also recognized to be duplicated. However, *TraesCSU02G235300* showed duplications with genes located on chromosome 3B only, strongly suggesting that it was also located on the 3B chromosome. The MIKC-type MADS-box genes were also found to be more duplicated than *M*-type genes (60% vs 40% of duplicated gene pairs, respectively), particularly due to expended subfamilies (e.g. *AGL17, Bsister, MIKC^*^, SEP*, and *SOC1*). Remarkably, duplicated gene pairs were subfamily-specific and particularly recognized in subfamilies containing unbalanced homeologs, except for the *AP3* subfamily (Tables 1, **S5** & **S6**). Among subfamilies, *Mα, AGL17, MIKC^*^*, and *SEP* contained 25%, 21%, 14%, and 12% of the total duplicated gene pairs, respectively, and the majority of these were located in distal telomeric and sub-telomeric (one gene located on the telomeric segment and other on the central segment) chromosomal regions. We also observed that the majority of the duplicated gene pairs (~51%) were located in distal telomeric segments, whereas only 26% and 23% of the duplicated gene pairs were located in proximal and sub-telomeric segments of chromosomes. These results could be explained by higher gene density in distal vs central chromosomal segments (IWGSC, 2018). Furthermore, segmental duplications were more prevalent than tandem duplications (61% and 37%) in 197 of the duplicated gene pairs with available chromosomal information (**Table S6**). Interestingly, >37% of all tandem duplications were identified on chromosome 3B, consistent with IWGSC findings (IWGSC, 2018). Collectively, these results strongly suggest that unbalanced homeologs in distal telomeric regions derive MADS-box subfamilies expansion through segmental duplications. These results are supported by previous observations that distal chromosomal regions are frequent targets of recombination events and are hotspots for faster evolution of underlying genes (N. W. G. Chen et al., 2018; Glover et al., 2015; Ramírez-González et al., 2018).

**Fig. 3.**
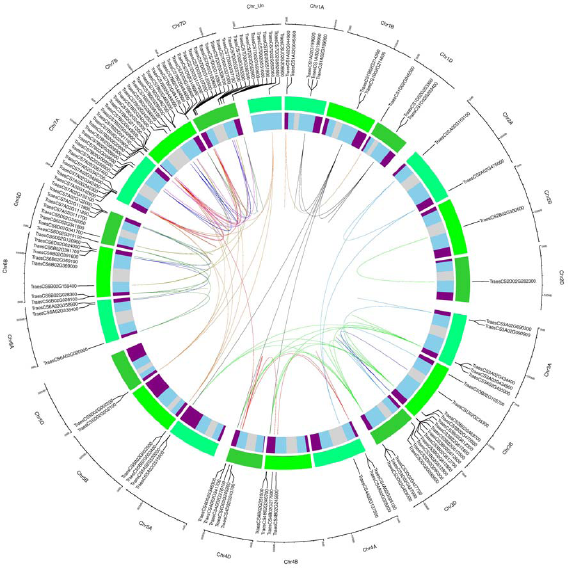
Subfamily-specific gene duplications among wheat MADS-box genes. Duplicated genes were plotted in a circular diagram using respective physical positions with shinyCircos (Yu et al., 2018). The outer track indicates three different subgenomes (shades of green) and inner track represents chromosomal segments (R and R3, purple; R2a and R2b, sky blue; C, light grey) (IWGSC, 2018). Duplicated genes were identified through sequence similarity (**Table S6**; see material and method section) and linked with subfamily-specific colours as in Fig. 1A & 1B, except for *AGL17*-like genes which were linked using chocolate colour. The linked duplicated genes positioned on different wheat chromosomes represent segmental duplications, whereas tandem duplications were indicated by incomplete links within the same chromosomes.

**Table 1.** Potential relationship between unbalanced homeologs and gene duplications in wheat MADS-box gene family.

| **Subfamily** | **Total Homeologs <sup>1</sup>** | **Balanced Homeologs <sup>2</sup>** | **Unbalanced Homeologs <sup>3</sup>** | **Gene Duplications** |
| --- | --- | --- | --- | --- |
| *Mα* | 53 | 33 | 20 | Present |
| *Mβ* | 28 | 10 | 18 | Present |
| *Mγ* | 47 | 26 | 21 | Present |
| *AG/STK* | 12 | 12 | 0 | Absent |
| *AGL12* | 6 | 6 | 0 | Absent |
| *AGL17* | 31 | 22 | 9 | Present |
| *AGL6* | 3 | 3 | 0 | Absent |
| *AP1* | 9 | 9 | 0 | Absent |
| *AP3* | 6 | 6 | 0 | Present <sup>4</sup> |
| *Bsister* | 19 | 10 | 9 | Present |
| *MIKC\** | 27 | 21 | 6 | Present |
| *OsMADS32* | 3 | 3 | 0 | Absent |
| *PI* | 6 | 6 | 0 | Absent |
| *SEP* | 28 | 25 | 3 | Present |
| *SOC1* | 13 | 12 | 1 | Present |
| *SVP* | 9 | 9 | 0 | Absent |

<sup>1</sup>, Total number of gene copies from three different wheat genomes (A, B, D); <sup>2</sup>, At least one gene copy is present in all diploid genomes; <sup>3</sup>, Gene copy is missing in one or two genomes;
<sup>4</sup>, Exception-gene duplications identified despite holding balanced homeologs.

### 2.5. Faster evolution of M-type MADS-box genes

To investigate evolution rates, we estimated substitutions in coding sequences and computed approximate divergence time between duplicated gene pairs (**Table S6**). In M-type genes, 1^st^ and 3^rd^ quartiles of synonymous substitutions (Ks) were considerably narrower than MIKC-type genes, whereas non-synonymous to synonymous substitution ratios (Ka/Ks) of M-type genes were significantly higher than 1^st^ and 3^rd^ quartiles of MIKC-type genes (Fig. 4). Furthermore, the percentage of duplication events as a result of positive/Darwinian selection (Ka/Ks ration > 1) was almost doubled (23%) in M-type genes as compared with MIKC-type genes (12%) (**Table S6**). Additionally, the estimated divergence time of M-type genes was significantly narrower than the MIKC-type genes. The mean divergence time of M-type genes was nearly half of the mean of MIKC-type genes (Fig. 4). Taken together, these data strongly indicate a more recent, rapid evolution of M-type MADS-box genes, which could be explained by their predominant localization in distal telomeric regions of chromosomes (Fig. S3 & S4; **Table S6**).

**Fig. 4.**
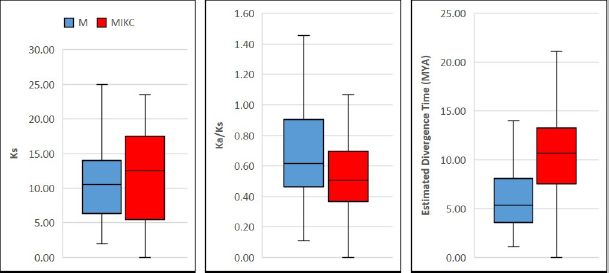
Box-plots showing evolution patterns in M- and MIKC-type wheat MADS-box genes. The coding sequences of the duplicated genes were MAFFT aligned (Katoh et al., 2018; Katoh and Standley, 2013), masked and subjected to SNAP V2.1.1 program for computing the synonymous (Ks) and non-synonymous (Ka) substitution rates. The divergence time between duplicated genes pairs was estimated in million years ago (MYA) by following El Baidouri et al. (2017) and box-plots were generated using Microsoft Excel 2016.

### 2.6. Expression patterns of MADS-box genes under abiotic and biotic stresses

Extensive investigations have been carried out to study the expression patterns of MADS-box genes during growth and development of crop plants. However, their transcriptional regulation under stressful conditions is somewhat obscure. Therefore, we analyzed RNA-seq based expression data of 300 MADS-box genes under all available biotic and abiotic stress conditions in the exVIP wheat expression browser (Borrill et al., 2016; Ramírez-González et al., 2018) (Fig. 5; **Table S7**). Out of the total 300 genes, nearly 57% were expressed (log_2_ TPM 0.20–7.89) during at least one developmental stage of one or more of the stresses included in this study. Whereas, the remaining 43% of genes showed no or very low expression (log_2_ TPM < 0.0), and were subsequently considered as not expressed.

**Fig. 5.**
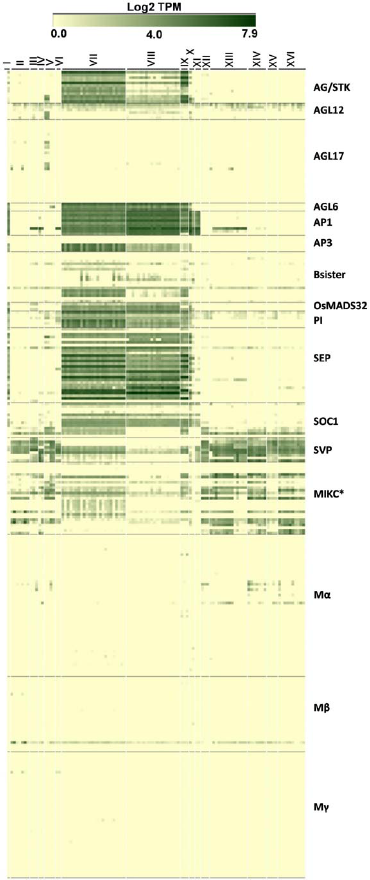
Expression patterns of wheat MADS-box genes under stressful conditions. RNA-seq based expression data of all MADS-box genes were retrieved from exVIP wheat expression browser (Borrill et al., 2016; Ramírez-González et al., 2018) and a heatmap was generated with Genesis software (Sturn et al., 2002). Processed expression levels of all genes under different abiotic (I, spikes with water stress; II, drought & heat-stressed seedlings; III, seedling treated with PEG to induce drought; IV, shoots after two weeks of cold stress; V, phosphate starvation in roots/shoots/leaves) and biotic stresses (VI, coleoptile infection with *Fusarium pseudograminearum*/crown root; VII, FHB infected spikelets (0 to 48 hours); VIII, FHB infected spikelets (30 to 50 hours); IX, spikes inoculated with FHB and ABA/GA; X, CS spikes inoculated with FHB; XI, leaves naturally infected with *Magnaporthe oryzae*; XII, PAMP inoculation of seedlings; XIII, stripe rust infected seedlings; XIV, stripe rust and powdery mildew infection in seedlings; XV, *Septoria tritici* infected seedlings; XVI, *Zymoseptoria tritici* infected seedlings) (columns) and in different subfamilies (rows) are presented as Log_2_ transcripts per million (Log_2_ TPM). Detailed information about expression levels of individual genes in tested tissues/growth stages and stresses/diseases are provided in **Table S7**.

Overall, MIKC-type MADS-box genes were highly expressed under all studied stresses, whereas nearly 75% of the non-expressed genes belonged to the three M-type subfamilies. Interestingly, the remaining 25% of the non-expressed genes belonged to MIKC-subfamilies containing duplicated genes, e.g. *AGL17, MIKC^*^*, and *Bsister*. As the majority of the duplicated genes were located in distal telomeric and sub-telomeric segments of chromosomes, it could be expected that promoters of these genes had undergone H3K27me3 hypermethylation which resulted in their lower or none existent expression (Ramírez-González et al., 2018).

In general, genes of all MIKC-type subfamilies (except *AGL17*) were ubiquitously expressed in fusarium head blight infected spikelets/spikes. Nevertheless, floral homeotic genes (*AP1, AP3, PI, AG/STK, AGL6*, and *SEP*) showed highest expression patterns. *SVP, MIKC^*^, AGL12*, and *SOC1*-like genes were also ubiquitously expressed in all studied abiotic and biotic stresses, strongly indicating that genes belonging to these subfamilies are active guardians of the wheat genome under stressful environments. Remarkably, all *AP1*-like genes demonstrated high expression in *Magnaporthe oryzae* infected leaves, suggesting disease-specific expression, as well as, the involvement of these important subfamily genes in diverse aspects of wheat growth and development. Similarly, *AGL17*-like genes were expressed only in seedlings under drought, heat, and phosphate starvation conditions.

We also calculated expression based hierarchical clusters to analyze diversification in expression patterns present within subfamilies (Fig. S5, **Table S8**). The *MIKC^*^* and *SOC1*-like genes were grouped into seven and six random clusters, respectively. Similarly, *Bsister* and *SEP*-like genes were separately grouped into five random clusters. By contrast, *AP1* and Mγ-like genes exhibited no variation in their expression patterns, as all genes were grouped into a single random cluster. Likewise, *AGL6, AGL17, AP3, OsMADS32, Mα*, and *Mβ*-like genes displayed little diversification in expression, and genes of each subfamily were grouped into two random clusters. In comparison, the majority of the genes with no or very low expression (~58%) were grouped into cluster 10 and all belonged to subfamilies with duplicated genes (Fig. S4, **Table S8**).

## 3. Discussions

### 3.1. Recent significant advancements in identification and characterization of MADS-box family members in wheat

MADS-box gene family members have been identified across all groups of eukaryotes and are known to confer diverse biological functions. In plants, these play a central role during growth and development, and are thus very important targets for crop improvement (Ali et al., 2019). Recently, significant advancements have been made in identification and characterization of MADS-box family members in wheat, which could in turn facilitate crop breeding efforts and aid in development of more resilient and higher-yielding cultivars. Ma et al. (2017) reported the first genome-wide analysis of MADS-box family members and identified a total of 180 MADS-box genes in an earlier genome version (IWGSC, 2014). Their *in-silico* expression data provided insights into the stress associated functions of MADS-box genes. To add to this, Schilling et al. (2020) conducted a comprehensive analysis of MIKC-type MADS-box genes and identified a total of 201 genes from an updated genome version (Ref. Seq 1.0) (IWGSC, 2018). They speculated that pervasive duplications, functional conservation, and putative neofunctionalization may have contributed to the adaptation of wheat to diverse environments. These advancements, along with results presented in this study, have begun to elucidate the broad landscape of wheat MADS-box genes and provide new insights into the phylogenomic, evolution, and functions of MADS-box family members.

### 3.2. MADS-box genes are underestimated in the wheat genome

Bread wheat is a hexaploid species (AABBDD, 2n=6x=42), which originated from a series of naturally occurring hybridization events among three closely related and independently maintained sub-genomes. Its full genome size is ~17 Gbps, among which ~14.5 Gbps (85%) is sequenced and contains ~107,891 coding genes (IWGSC, 2018). In this study, through a comprehensive genome-wide analysis, we identified a total of 300 high confidence, full length, non-redundant MADS-box genes from the latest publicly available reference genome version (Fig. 1, **Table S2**). To date, this is the second-highest number of MADS-box genes identified in a crop plant, closely following *Brassica napus* (Wu et al., 2018), which contains 307 full-lengths and/or incomplete (pseudo) MADS-box genes. Possible explanations for the abundance found in wheat may encompass the larger genome size, higher gene number, high rate of homeolog retention, and hexaploid nature of bread wheat (IWGSC, 2018). Moreover, at least 35 low confidence MIKC-type MADS-box genes were recently reported in the IWGSC reference genome (Schilling et al., 2020). If these low confidence genes are also included with the 300 MADS-box genes identified in this study, then the final number will be the highest identified in a plant genome. Furthermore, a more recent chromosome-scale assembly of the bread wheat genome revealed hundreds of new genes and thousands of additional gene copies (Alonge et al., 2020) which were substantially missing from the IWGSC reference genome explored in the current study. Collectively, all these observations strongly suggest that MADS-box genes are underestimated in wheat and their exploration might help in understanding the global adaptability of wheat.

### 3.3. Does grass genomes lack FLC-like genes?

*FLOWERING LOCUS C*s or *FLC*s are key genes conferring vernalization requirement, which act as flowering repressors in *Arabidopsis* (Whittaker and Dean, 2017). High expression of *FLCs* repress other flowering activator genes and causes delayed flowering. In winter varieties of temperate cereals, including barley, *Brachypodium*, and wheat, vernalization is regulated through *VERNALIZATION* genes (*VRN1, VRN2, VRN3*) (Greenup et al., 2009). To date, strong disagreement exists in the published literature regarding the presence or absence of *FLC*-like genes in cereals (Paolacci et al., 2007; Ruelens et al., 2013; Schilling et al., 2020; Wei et al., 2014; Zhao et al., 2011). In this study, despite using updated resources and tools, we were unable to find *FLC*-like genes within the wheat or rice genomes (Fig. 1A). Interestingly, wheat genes which were previously reported to be *FLC*-like (Ruelens et al., 2013; Schilling et al., 2020), as well as *OsMADS37* and *OsMADS51*/*65*, were grouped with *MIKC^*^*-like genes (Fig. 1A, **Table S4**). In comparison, all other *FLC* homologs were grouped into a separate distinct *Arabidopsis*-specific *FLC* clade. Moreover, we also demonstrated that multiple sequence amino acid alignments of *FLC* and *MIKC^*^*-like genes were significantly dissimilar and none of the grass genes shared sequence homology with *Arabidopsis FLC* paralogs (Fig. 2). Altogether, these results indicate that *FLC*-like genes might have been lost in cereal genomes after they diverged from eudicots. Several other studies are also in agreement with current results and were unable to identify *FLC*-like genes in grass genomes (Arora et al., 2007; Paolacci et al., 2007; Wei et al., 2014; Zhao et al., 2011). These observations suggest that vernalization pathways in monocots and dicots evolved independently and are regulated through a completely different set of genes. However, earlier reports on the identification of *FLC*-like genes in grasses (Ruelens et al., 2013; Schilling et al., 2020; Sharma et al., 2017) and functional conservation of *ODDSOC2* with *FLC* in the regulation of vernalization (Greenup et al., 2010), suggest that evolution of vernalization pathways between monocots and dicots may not be fully independent. Thus, the biological question of the presence or absence of *FLC*-like genes in grass genomes still needs to be explored. Further research efforts in this direction might address this fundamental question, and propitious outcomes could help in breeding efforts to develop winter varieties of temperate cereals which are adapted to changing environments.

### 3.4. Subfamily-specific distal telomeric duplications in unbalanced homeologs facilitate rapid adaptation to changing environments

Distal telomeric chromosomal segments are evolutionary hotspots for frequent recombination events and give rise to fast-evolving genes (N. W. G. Chen et al., 2018; Glover et al., 2015). Many adaptability trait related genes which are induced in response to external stimuli are found to be predominately located in distal chromosomal segments. In comparison, genes related to housekeeping and conserved developmental functions are positioned in central chromosomal segments (IWGSC, 2018; Ramírez-González et al., 2018). In this study, the majority of the identified duplicated gene pairs were located in distal telomeric (~51%) and sub-telomeric (23%) chromosomal segments (Fig. 3, **Table S6**). Remarkably, all these duplications were subfamily-specific and primarily identified in larger subfamilies containing unbalanced homeologs (*Mα, Mβ, Mγ, AGL17, Bsister, MIKC*, SEP*, and *SOC1*) (Table 1). Interestingly, genes belonging to these subfamilies have been reported to regulate plants adaptability in changing environments. For example, *OsMADS57* (*AGL17*-like gene in rice) is induced by abscisic acid, chilling, drought, and salinity stress (Arora et al., 2007), promotes cold tolerance by interacting with a defence gene (*OsWRKY94*) (L. Chen et al., 2018), and modulates root to shoot nitrate translocation under deprived nitrate conditions (Huang et al., 2019). Similarly, downregulation of a *SEP* clade gene in pepper (*CaMADS*) caused more sensitivity to cold, salt, and osmotic stresses, whereas its overexpression in *Arabidopsis* conferred higher tolerance against these stresses (Chen et al., 2019). By contrast, almost negligible duplications were identified in a single smaller subfamily (*AP3*) with completely balanced homeologs (Table 1). These results strongly indicate that unbalanced homeologs of expended subfamilies have undergone subfamily-specific distal telomeric duplications, thus facilitating the rapid adaptation of bread wheat to diverse global environments. Contrarily, completely balanced homeologs of functionally conserved smaller subfamilies might have some evolutionary advantage in minimizing the developmentally detrimental gene copy number variations, due to their localization in central chromosomal segments.

### 3.5. A recent and faster evolutionary origin of M-type MADS-box genes

Unlike MIKC-type MADS-box genes, the M-type genes have not been extensively studied and little information is available about their evolutionary origin in crop plants (Masiero et al., 2011). In this study, we observed that M-type genes were predominately located in distal telomeric chromosomal segments (**Table S2**) and are evolving at a faster rate, probably due to higher frequency of tandem gene duplications and stronger purifying selections (Fig. 3, **Table S6**). Moreover, lesser synonymous substitutions and higher non-synonymous to synonymous substitution ratios of M-type duplicated genes exposed their recent divergence and evolutionary origin as compared with MIKC-type genes (Fig. 4, **Table S6**). These results are in agreement with a study by Nam et al. (2004), in which they also reported faster birth-and-death evolution of M-type genes in angiosperms. This data might indicate the significance of M-type MADS-box genes in understanding the evolutionary history of the bread wheat genome.

### 3.6. MADS-box genes as active guardians of plants against pathogen insurgency and harsh environments

MADS-box genes regulate diverse developmental processes and their functions are well studied in plant morpho- and organogenesis (Ali et al., 2019; Smaczniak et al., 2012). However, several members of the MADS-box gene family are reported to be involved in the regulation of biotic and abiotic stress responses (Castelán-Muñoz et al., 2019; Q. Wang et al., 2018; Zhang et al., 2016), which point towards possible dynamic roles in stress response. In this study, we analyzed the complete atlas of their expression profiles in all publicly available wheat transcriptomic data under biotic and abiotic stresses (Fig. 5 & S5; **Tables S7** & **S8**). More than 50% of the total identified MADS-box genes were expressed during at least one developmental stage of one or more stresses, and the majority of these expressed genes belonged to the MIKC-group (**Table S7**). Except for *AGL17*, genes of all other MIKC-type subfamilies were ubiquitously expressed in FHB infected spikes/spikelets (Fig. 5 & S5), mimicking their defensive roles against *Fusarium* infection. Previously, Yang et al. (2015) functionally characterized *FgMcm1* in the causal agent (*Fusarium graminearum*) of barley and wheat head blight disease and demonstrated that *FgMcm1* played crucial roles in cell identity and fungal development. More recently, Xu et al. (2020) also reported a close relationship between anther extrusion and field FHB resistance and pointed out that *Rht* and *Vrn* genes might have pleiotropic effects on these traits. Since FHB is a floral disease and floral organ identity is controlled by MIKC-type MADS-box genes (Ali et al., 2019), this indicates a cooperate relationship for their involvement in the pathogen response network. In future, it would be interesting to investigate how MADS-box genes are involved in fighting FHB infection.

Similarly, *SVP, MIKC^*^, AGL12*, and *SOC1* subfamily genes were ubiquitously expressed under all studied stresses (Fig. 5, **Table S7**), pointing towards their more critical role in stress associated functions. In comparison, members of the largest MIKC group (*AGL17*) were expressed under phosphate starvation, and heat and drought stress conditions. Several functionally characterized genes belonging to all these subfamilies have also been reported to regulate different abiotic and/or biotic stresses in plants (Hwang et al., 2019; Khong et al., 2015; P. Li et al., 2020; SHI et al., 2016; Z. Wang et al., 2018). Interestingly, few M-type genes, especially *Mα*-like, were weakly expressed under different biotic stresses (Fig. 5) and these expression patterns are in agreement with a Guo et al. (2013) study in which they observed differential expression of an M-type gene in response to stripe rust infection in wheat. Collectively, these results strongly highlight MADS-box genes, especially MIKC-type, as key members of gene regulatory networks (Castelán-Muñoz et al., 2019) and their involvement in wheat response to possible pathogen insurgency and harsh environments.

### 3.7. Towards bridging the gap between genotype and phenotype using genetic and genomic resources

Understanding the functions of candidate genes controlling agronomically important traits is critical for the acceleration of crop improvement efforts. Following the unveiling of recent wheat genetic and genomic resources (Adamski et al., 2020; Avni et al., 2017; Luo et al., 2017; Maccaferri et al., 2019), rapid testing of previous model plant discoveries and their cross-application in staple crops is paving the way towards bridging the gap between genotype and phenotype. During recent years, several wheat genes related to flowering, yield, quality, disease resistance, male sterility, and nutrient use efficiency traits have been cloned and functionally characterized (briefly reviewed in Li et al. (2020)), which exposed the unprecedented potential of these genetic and functional genomic resources in establishing genotype-to-phenotype relationships. These resources could also facilitate in elucidating the regulatory roles of neglected MADS-box subfamilies, such as *Mα, Mβ, Mγ*, and *MIKC^*^*, during wheat growth and development, and subsequently help in deciphering whether MADS-box genes confer pathogen resistance and/or abiotic stress tolerance. Furthermore, many other fundamental biological questions about their phylogenomics, evolutionary origin, and stress associated functions, could be addressed by exploiting these paramount resources.

In conclusion, MADS-box genes are critically important for wheat growth and development, and hold enormous promise for bolstering yield potential under changing global environments. Our results speculate that the abundance of MADS-box genes in the wheat genome might be associated with its global adaptability and that subfamily-specific distal telomeric duplications in unbalanced homeologs facilitate its rapid adaptation. In addition, through comprehensive genome-wide and comparative analyses, we demonstrated that wheat and rice genomes lack *FLC* genes, which could help molecular breeders in the identification of target genes for fine-tuning of winter wheat varieties. Moreover, our *in-silico* expression data strongly indicated the possibility of protective roles of MADS-box genes against pathogen attacks and harsh climatic conditions. In this way, we provided an entire complement of MADS-box genes identified in the wheat genome that will accelerate functional genomics efforts and possibly facilitate in bridging genotype-to-phenotype relationships through fine-tuning of agronomically important traits.

## 4. Material and methods

### 4.1. Identification of MADS-box genes

The latest publicly available genome version of wheat (IWGSC Ref. Seq v1.1) was accessed through the ensemble plants database (Bolser et al., 2017) and searched using protein family database (Pfam) identifiers of MADS (PF00319) and K (PF01486) domains. A total of 281 MADS-domain and 131 K-domain encoding genes were identified. Additionally, a query-based search using “MADS” yielded 300 genes. Together, these searches identified a total of 712 genes (**Table S1**), among which 300 were found to be high confidence and non-redundant and considered for further analyses. Detailed information of these 300 genes was retrieved from the ensemble plants database and is provided in **Table S2**. Moreover, amino acid sequences for the identified genes were uploaded into the NCBI conserved domain database (Marchler-Bauer et al., 2015) for validation of putative protein domains (**Table S3**). Gene identifiers were used as gene names for subsequent analyses; however, alternative names used in previous studies (Schilling et al., 2020) were also provided in **Table S4**.

### 4.2. Comparative phylogenetic analyses and subfamily classification

Differentiation between type I (M-type) and type II (MIKC-type) MADS-box proteins were achieved by separate MAFFT alignments using only the MADS domain (L-INS-i algorithm) (Katoh et al., 2018; Katoh and Standley, 2013) between *Arabidopsis* (Pařenicová et al., 2003) and wheat, and rice (Arora et al., 2007) and wheat MADS-box proteins. Subsequently, maximum likelihood (ML) phylogenies were inferred using I_Q_-T_REE_ (Nguyen et al., 2015) by choosing JTT+F+G4 best fit substitution model according to the Bayesian information criterion (BIC) (Kalyaanamoorthy et al., 2017). Consistency of the ML trees was validated by setting an Ultrafast bootstrap value of 1000 (Hoang et al., 2018; Minh et al., 2013). The final phylogenetic trees were visualized with MEGA7 (Kumar et al., 2016).

Subfamily classifications were accomplished by separate MAFFT alignments within M-type (L-INS-i algorithm) and MIKC-type (E-INS-i algorithm) MADS-box protein sequences of *Arabidopsis*, rice, and wheat (Katoh et al., 2018; Katoh and Standley, 2013). The full-length alignments were subjected to the Gap Strip/Squeeze v2.1.0 tool (www.hiv.lanl.gov/content/sequence/GAPSTREEZE/gap.html) for masking the individual residues by removing the gaps with default parameters. Then, masked alignments of M-type and MIKC-type proteins were independently subjected to the I_Q_-T_REE_ software (Nguyen et al., 2015) for generating ML phylogenetic trees as described above. Subfamily names were given by following subfamily classifications in *Arabidopsis* and/or major grass species (Gramzow and Theißen, 2015) (**Table S2** & **S4**).

### 4.3. Homeologs identification

Putative homeologs were recognized based on strong phylogenetic relationships (Ultrafast bootstrap value > 90) within different sub-families. Classifications reported in previous studies were also considered (IWGSC, 2018). The homeologs status of thirty-one genes could not be determined due to lower Ultrafast bootstrap values (**Table S5**).

### 4.4. Gene duplication and evolution analyses

Coding sequences (CDS) of all wheat MADS-box genes were retrieved from the ensemble plants database and blasted against each other using Sequence Demarcation Tool V1.2 (Muhire et al., 2014) for the identification of sequence identities. Gene pairs with ≥ 90% identity (E value <1e^-10^) and non-homeologous status were considered as duplicated (Ning et al., 2017) (**Table S6**). If the duplicated homologous gene pair was located on the same chromosome it was defined as tandem duplication. Otherwise, when homologous gene pairs were located on different chromosomes it was defined as segmental duplication. The CDS sequences of duplicated genes were MAFFT aligned, masked, and subjected to the Synonymous Non-Synonymous Analysis Program V2.1.1 (www.hiv.lanl.gov/content/sequence/SNAP/SNAP.html) to compute the synonymous (Ks) and non-synonymous (Ka) substitution rates. To find out which type of codon selection operated during evolution, the ratio of Ka/Ks was also calculated. The approximate divergence time between duplicated gene pairs was calculated by using the formulae T=Ks/2r × 10^-6^ assuming a substitution rate (r) of 6.5 × 10^-9^ substitutions/synonymous site/year (El Baidouri et al., 2017) (**Table S6**).

### 4.5. In-silico expression analyses

Expression data under all available abiotic and biotic stress conditions were retrieved from the expVIP Wheat Expression Browser (Borrill et al., 2016; Ramírez-González et al., 2018) as Log_2_ TPM (processed expression value in transcripts per million) obtained via RNA-seq analysis. Detailed information about expression levels of individual genes in tested tissues/growth stages and stresses/diseases are provided in the supplementary information (**Table S7**). Genesis software (Sturn et al., 2002) was used to generate heatmaps from the obtained expression data. Expression based clustering of genes was achieved by following the *K*-means clustering method (K = 10, iterations = 1000, runs = 5) (**Table S8**).

## Acknowledgments

All authors are grateful to Dr Yuling Jiao, Institute of Genetics and Developmental Biology, Chinese Academy of Sciences for his comments and suggestions on the manuscript. The inspiration and support from Precision Agriculture and Analytics Lab (PAAL), affiliate of National Center in Big Data and Cloud Computing (NCBC) is also gratefully acknowledged.

## Author contributions

**Qasim Raza:** Conceptualization, Data curation, Formal Analysis, Methodology, Software, Validation, Visualization, Writing-original draft. **Awais Riaz:** Data curation, Formal Analysis, Methodology, Software, Validation, Visualization, Writing-review & editing. **Rana Muhammad Atif:** Data curation, Supervision, Validation, Writing-review & editing. **Babar Hussain:** Data curation, Visualization, Writing-original draft. **Zulfiqar Ali:** Conceptualization, Supervision, Writing-review & editing. **Hikmet Budak:** Supervision, Writing-review & editing.

---

**Fig. S1.**
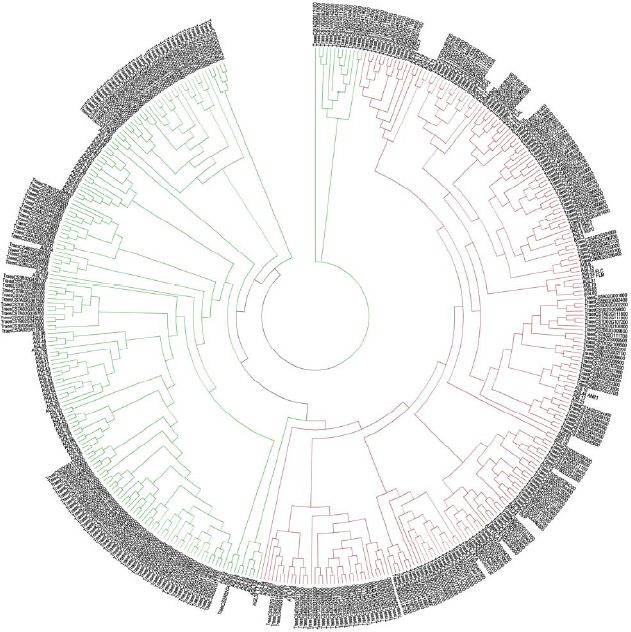
Comparative phylogenetic tree between *Arabidopsis* and wheat MADS-box genes. *Arabidopsis* and wheat MADS-box proteins were MAFFT aligned using only MADS domain (L-INS-i algorithm), maximum likelihood phylogenies inferred using I_Q_-T_REE_ software and final tree visualized with MEGA7. M- and MIKC-type genes were highlighted with green and red colour nodes, respectively.

**Fig. S2.**
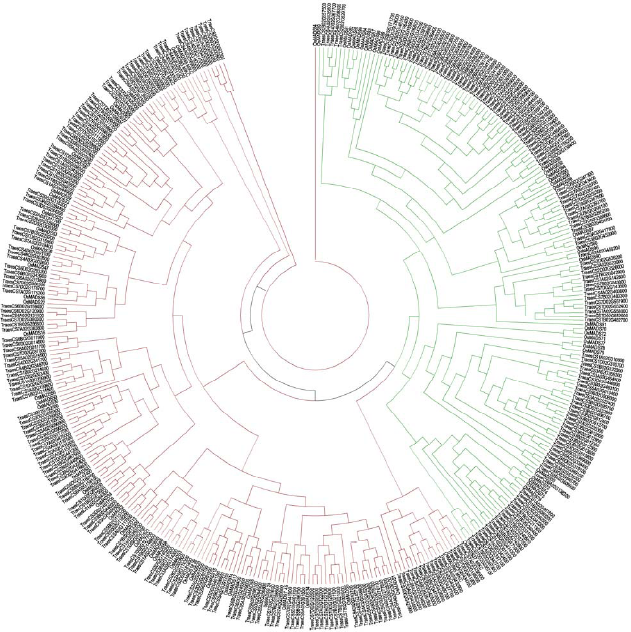
Comparative phylogenetic tree between rice and wheat MADS-box genes. Rice and wheat MADS-box proteins were MAFFT aligned using only MADS domain (L-INS-i algorithm), maximum likelihood phylogenies inferred using I_Q_-T_REE_ software and final tree visualized with MEGA7. Mand MIKC-type genes were highlighted with green and red colour nodes, respectively.

**Fig. S3.**
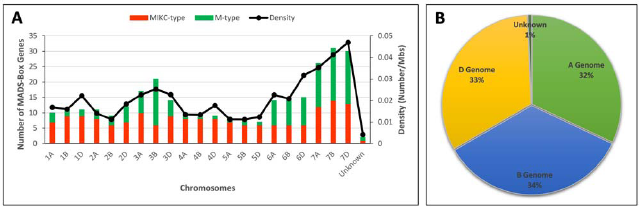
Distribution pattern of MADS-box genes on wheat chromosomes and sub-genomes.

**Fig. S4.**
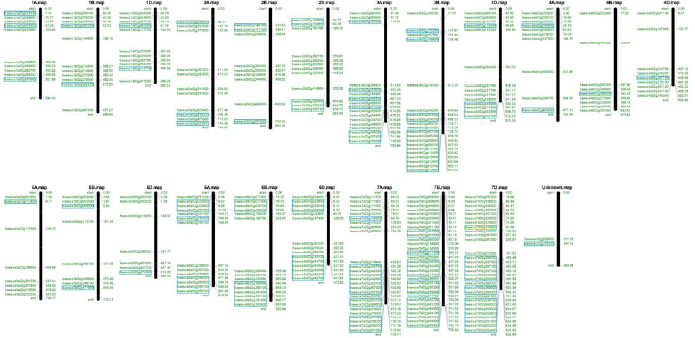
Chromosomal mapping of wheat MADS-box genes. Solid black vertical bars represent individual chromosomal lengths (Mbp). Gene names and their initial physical positions are drawn on the left and right side of the respective chromosomes, respectively. M-type genes are encircled with blue boxes.

**Fig. S5.**
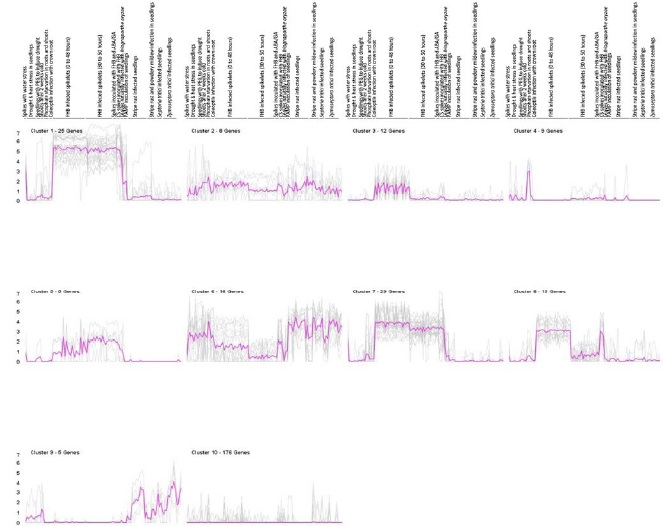
Expression patterns based clustering of wheat MADS-box genes. Expression patterns based clustering of genes was achieved by following the K-means clustering method (K = 10, iterations = 1000, runs = 5) with Genesis software (**Table S8**).

